# Sterile triggers drive joint inflammation in TNF and IL-1β dependent mouse arthritis models

**DOI:** 10.1101/2023.01.31.526410

**Authors:** Alexandra Thiran, Ioanna Petta, Gillian Blancke, Marie Thorp, Guillaume Plankaert, Maude Jans, Vanessa Andries, Korneel Barbry, Tino Hochepied, Christian Vanhove, Eric Gracey, Emilie Dumas, Teddy Manuelo, Ivan Josipovic, Geert van Loo, Dirk Elewaut, Lars Vereecke

## Abstract

Arthritis is the most common extra-intestinal complication in inflammatory bowel disease (IBD). Conversely, arthritis patients are at risk for developing IBD and often display subclinical gut inflammation. These observations suggest a shared disease etiology, commonly termed ‘the gut-joint-axis’. The clinical association between gut and joint inflammation is further supported by the success of common therapeutic strategies and microbiota dysbiosis in both conditions. Most data however support a correlative relationship between gut & joint inflammation, while causative evidence is lacking. Using two independent transgenic mouse arthritis models, either TNF or IL1β dependent, we demonstrate that arthritis develops independently of the microbiota and intestinal inflammation, since both lines develop full-blown articular inflammation under germ-free conditions. In contrast, TNF-driven gut inflammation is fully rescued in germ-free conditions indicating that the microbiota is driving TNF-induced gut inflammation. Together, our study demonstrates that, although common inflammatory pathways may drive both gut and joint inflammation, the molecular triggers initiating such pathways are distinct in these tissues.

## Introduction

There is convincing evidence that gut and joint inflammation are clinically linked, particularly in spondyloarthritis (SpA), a group of inflammatory joint diseases which can affect both peripheral joints and the spine, and often present with extra-articular manifestations including ileitis, colitis, psoriasis and uveitis (Taurog J, Chhabra A, 2016). Interestingly, acute infections with enteric pathogens like *Salmonella, Shigella* and *Campylobacter* can trigger reactive arthritis (ReA) (Taurog J, Chhabra A, 2016). 50% of all SpA patients present with subclinical gut inflammation, diagnosed by histological presence of microscopic gut inflammation, and 10% of all SpA patients eventually develop inflammatory bowel disease (IBD) (Leirisalo-Repo *et al*., 1994; Mielants, Veys, Cuvelier, De Vos, Goemaere, De Clercq, Schatteman and Elewaut, 1995; Mielants, Veys, Cuvelier, De Vos, Goemaere, De Clercq, Schatteman, Gyselbrecht, *et al*., 1995; Mielants, Veys, De Vos, *et al*., 1995; Van Praet *et al*., 2013). Genome wide association studies (GWAS) have revealed several shared disease susceptibility loci in IBD and SpA, including genes associated to innate immunity, type 3 immunity, and intestinal barrier integrity (Gracey *et al*., 2020). Various arthritic diseases, including rheumatoid arthritis (RA) and SpA, are characterized by shifts in intestinal microbial community structure and composition, and while this dysbiosis sometimes precedes arthritic disease onset, it remains unclear whether and to what extent it causally contributes to arthritis development (Breban *et al*., 2017; Ciccia *et al*., 2017; Tito *et al*., 2017; Zaiss *et al*., 2021). These observations have led to the dogma that joint inflammation is triggered or modulated by microbial signals and intestinal inflammation, and thus that intestinal pathology may precede and instigate joint inflammation. Various hypotheses have been suggested to support this idea, including the ‘arthritogenic peptide hypothesis’ which suggests that microbial derived antigens induce autoreactive T cells trough molecular mimicry, the ‘aberrant trafficking hypothesis’ which claims that immune cells primed in the intestinal mucosa home to synovial tissues and cause inflammation, and the ‘dysbiosis hypothesis’ which states that a shift in microbiota composition drives both intestinal and joint inflammation through various mechanisms (Qaiyum, Lim and Inman, 2021). Despite the strong correlation of gut and joint inflammation in SpA, there is no consensus that inflammation in the joint depends on intestinal dysbiosis or inflammatory events in the gut. Shared inflammatory pathways may underly both gut and joint inflammation, but can be triggered by independent and tissue-specific factors, which can either be microbial-derived or sterile triggers. Previous studies have demonstrated that TNF-driven intestinal inflammation in TNF^ΔARE^ mice (Kontoyiannis *et al*., 1999) is microbiota dependent, as TNF^ΔARE^ mice only develop spontaneous ileitis in colonized conditions, but not when raised under germ-free (GF) conditions (Roulis *et al*., 2016; Schaubeck *et al*., 2016). Despite the clear protection from intestinal inflammation, it is not clear whether GF TNF^ΔARE^ mice are equally protected from spontaneous joint inflammation. In order to investigate the gut-joint axis in detail, we studied two transgenic mouse models of arthritis, one which depends on the cytokine TNF, and one which is IL-1β driven, and evaluated the development of gut and joint inflammation in both colonized (specific pathogen free, SPF) and in GF conditions.

## Results

### Development and characterization a new TNF-driven transgenic mouse model

Given the importance of TNF in various human inflammatory diseases, including IBD and SpA, we generated a new TNF-driven mouse inflammation model by targeting the AU-Rich element (ARE) of the *Tnf* gene using a double guide-RNA mediated CRISPR/CAS9 approach, resulting in a 107 bp deletion in the 3’ UTR of the *Tnf* gene on Chromosome 17 (Figure S1). This deletion is predicted to generate a more stable *Tnf* mRNA compared to wild-type *Tnf* RNA, since binding of the ARE sequence and subsequent mRNA decay by TIS11 family RNA-binding proteins is prevented. The more stable mRNA results in elevated levels of bioactive TNF protein upon translation (Kontoyiannis *et al*., 1999; Makita, Takatori and Nakajima, 2021). This new TNF overexpressing mouse line, C57Bl6/J-Tnf^emARE1Irc^ (endonuclease modified, from now on termed **Tnf^emARE^**), was generated under SPF conditions and later rederived in GF conditions in the GF and gnotobiotic mouse facility at Ghent University. Macroscopically, both homozygous TNF^emARE/ARE^ and heterozygous TNF^emARE/+^ mice show stunted growth, in contrast to wild-type littermates (Figure 1A,B). Serum TNF levels are slightly elevated in heterozygous TNF^emARE/+^ mice, and significantly higher in homozygous TNF^emARE/ARE^ mice compared to wild-type littermates, confirming TNF overexpression in this model (Figure 1C).

**Figure 1:**
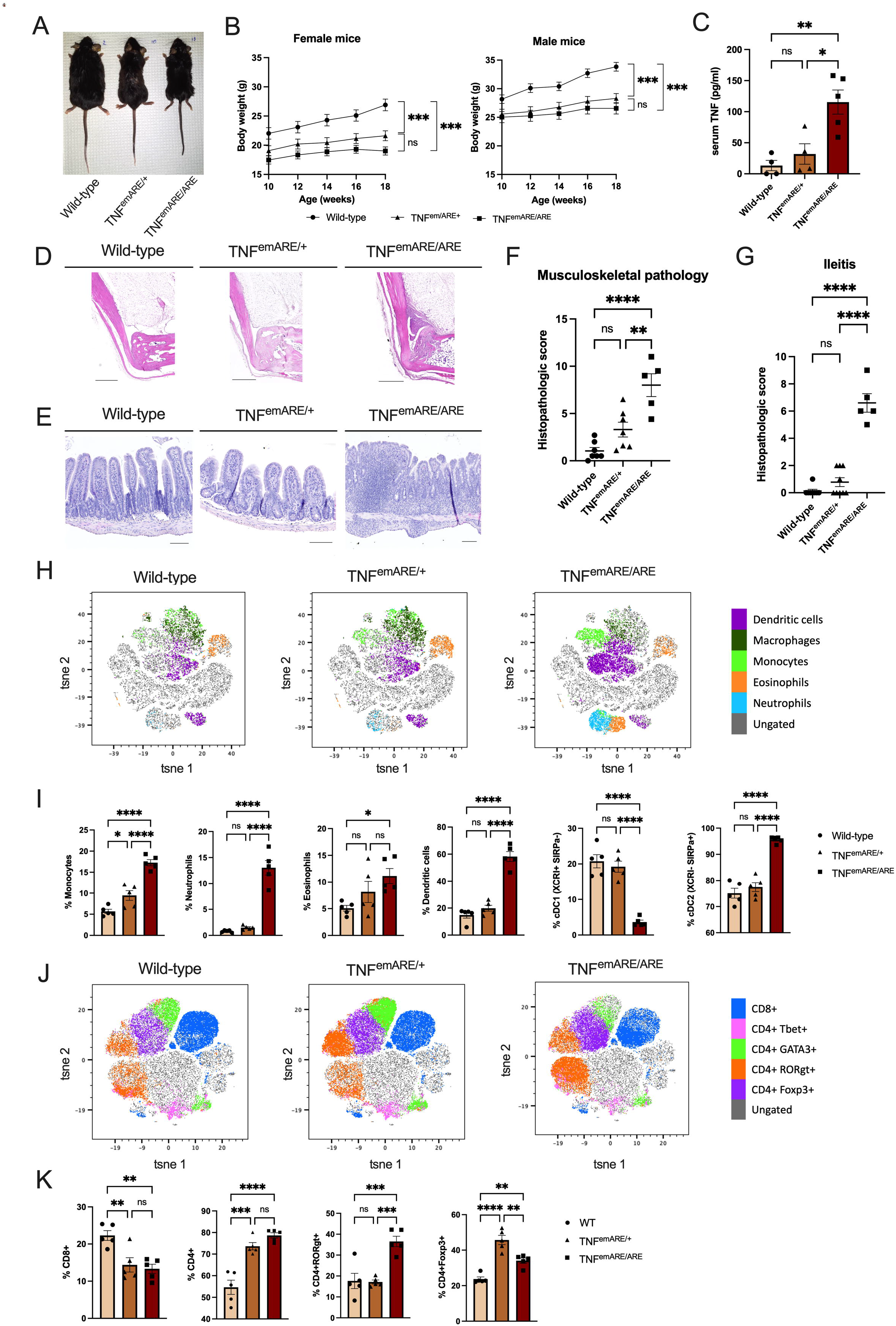
SPF TNF^emARE^ mice suffer from inflammatory gut and joint disease. (A) Macroscopic picture of a male wild-type, male TNF^emARE/+^ and male TNF^emARE/ARE^ mouse showing stunted growth in the hetero- and homozygous condition. Mice were 27-30 w/o. (B) Changes in body weight of wild-type (n=8; F=3, M=5), TNF^emARE/+^ (n=8; F=4, M=4) and TNF^emARE/ARE^ (n=8; F=6, M=3) mice over time for both male and female population. SPF TNF^emARE/ARE^ and TNF^emARE/+^ mice are smaller compared to wild-type littermates, which is reflected in their smaller body weight and limited body weight increase. (C) TNF^emARE/ARE^ mice show significantly elevated serum levels of TNF (wild-type n=4, TNF^emARE/+^ n=4, TNF^emARE/ARE^ n=5). (D-E) Histologic H&E sections of ileum and hind paw indicate severe ileitis and arthritis in 20-30 w/o SPF TNF^emARE/ARE^ mice (Scale bars upper panel: 200 μm, scale bars lower panel: 500 μm). (F) Quantification of musculoskeletal inflammation in hind paws of wild-type (n=7), TNF^emARE/+^ (n=9) and TNF^emARE/ARE^ (n=5) mice of 10-20 w/o. (G) Quantification of ileitis in wild-type (n=8), TNF^emARE/+^ (n=9) and TNF^emARE/ARE^ (n=5) mice of 10-20 w/o. (H) tSNE analysis performed on CD45+ lineage (excluding CD3+, CD19+, NK1.1+ fractions) of lamina propria of ileum samples (n=5/genotype). (I) Ileal flow cytometry data of 25 w/o TNF^emARE^ mice show a highly activated innate immune system (n=5/genotype). (J) tSNE analysis performed on CD3+ cells of lamina propria of ileum samples (n=5/genotype). (K) Ileal flow cytometry data show an increase of CD4+ cells in transgenic mice, which can be designated to the enrichment of the CD4+RORgt+ (Th17) cell population (n=5/genotype). (Plots are represented as Mean +/- SEM)

Peripheral musculoskeletal pathology was assessed by histological analysis of haematoxylin-eosin (H&E) stained ankle sections, demonstrating severe inflammation in TNF^emARE/ARE^ mice, characterized by immune cell infiltration along the Achilles tendon, in the synovial-entheseal complex (SEC) and Kargers’ fat pad and in the calcaneus (Figure 1D). The calcaneus shows bone erosion and mild bone marrow edema, a sign of systemic inflammation. In contrast, TNF^emARE/+^ mice develop mild to no obvious signs of peripheral and axial joint inflammation. Histopathological scoring based on signs of inflammation in the calcaneo-cuboid joint, the calcaneus and the synovium-entheseal complex (SEC) quantitatively supports our findings (Figure 1F). Gut pathology was also evaluated by histological analysis of H&E-stained ileal sections and revealed strong inflammation in TNF^emARE/ARE^ mice, which was minimal in heterozygous TNF^emARE/+^ mice and absent in wild-type littermate controls. Small intestinal inflammation in TNF^emARE/ARE^ mice is most severe in ileum, modest in jejunum and minimal in duodenum (Figure S2A), and is characterized by massive immune cell infiltration, loss of goblet cells and villus atrophy (Figure 1E, 1G). In contrast, we could not observe colonic inflammation in TNF^emARE/ARE^ mice (Figure S2B). Quantification of ileal lamina propria leukocytes by flow cytometry shows strongly increased populations of monocytes, eosinophils, neutrophils and dendritic cells in homozygous TNF^emARE/ARE^ mice (Figure 1H, 1I). Increased dendritic cell populations are primarily cDC2’s (XCRI-SIRPa+), while the cDC1 (XCRI+SIRPa-) population is drastically reduced in TNF^emARE/ARE^ mice. We also observe changes in the T cell compartment, including increased CD4+ T cell numbers and particularly CD4+RORgt cells (Figure 1 J,K). In contrast, in heterozygous TNF^emARE/+^ mice, ileal inflammation, and myeloid and T cell expansion is minimal. We confirmed that the inflammatory phenotype of TNF^emARE/ARE^ mice is dependent on TNF, since TNF^emARE/ARE^ mice are fully protected from spontaneous gut and joint inflammation when backcrossed in a TNF receptor 1 (TNFR1 or p55) deficient background (Figure S3).

In conclusion, this new TNF^emARE^ mouse line, which is characterized by TNFR1-mediated spontaneous Crohn’s-like ileitis, and peripheral and axial arthritis development resembles previously generated TNF^ΔARE^ mice (Kontoyiannis *et al*., 1999). In contrast to TNF^ΔARE^ mice, TNF^emARE^ mice only develop severe inflammatory pathology in homozygous conditions (TNF^emARE/ARE^ mice), while heterozygous TNF^emARE/+^ mice display minimal to no inflammation in gut and joints. We used this new TNF^emARE^ line to study microbiota dependency for the development of intestinal and joint inflammation.

### The microbiota instigates TNF-driven ileitis but not arthritis

Previous studies have shown that TNF^ΔARE^ mice are characterized by microbial dysbiosis, and that intestinal inflammation is microbiota-dependent (Roulis *et al*., 2016; Schaubeck *et al*., 2016). To investigate the contribution of the microbiota to TNF-driven inflammatory pathology in both gut and joints, TNF^emARE^ mice were rederived in germ-free (GF) conditions by axenic embryo transfer in the GF and gnotobiotic mouse facility at Ghent University. In contrast to SPF raised mice, GF raised TNF^emARE/ARE^ and TNF^emARE/+^ mice have similar body weight as their wild-type littermates (Figure 2A, 2B). Both SPF and GF TNF^emARE/ARE^ mice have significantly elevated TNF serum levels compared to wild-type littermate mice (Figure 2C). Moreover, ileal *Tnf* expression is significantly increased in SPF raised as well as in GF raised TNF^emARE/ARE^ mice (Figure 2D). However, expression of *Tnf* in the ileum of homozygote GF mice is lower compared to *Tnf* expression in their SPF counterpart. As TNF-mediated Paneth cell depletion was previously reported (Roulis *et al*., 2016), we performed immunostaining of Paneth cells by anti-lysozyme staining on ileal sections and observed complete Paneth cell depletion in TNF^emARE/ARE^ mice raised in SPF but not under GF conditions (Figure 2E). Ileal inflammation was further evaluated by H&E staining, and unlike SPF raised TNF^emARE/ARE^ mice which develop severe ileitis in SPF conditions, GF raised TNF^emARE/ARE^ mice are completely protected from ileitis (Figure 2F), which is confirmed by pathophysiological scoring of epithelial damage, structural changes and immune cell infiltration (Figure 2G). Together, these data confirm previous findings in TNF^ΔARE^ mice (Roulis *et al*., 2016; Schaubeck *et al*., 2016) and indicate that TNF-driven intestinal inflammation is fully dependent on the intestinal microbiota.

**Figure 2:**
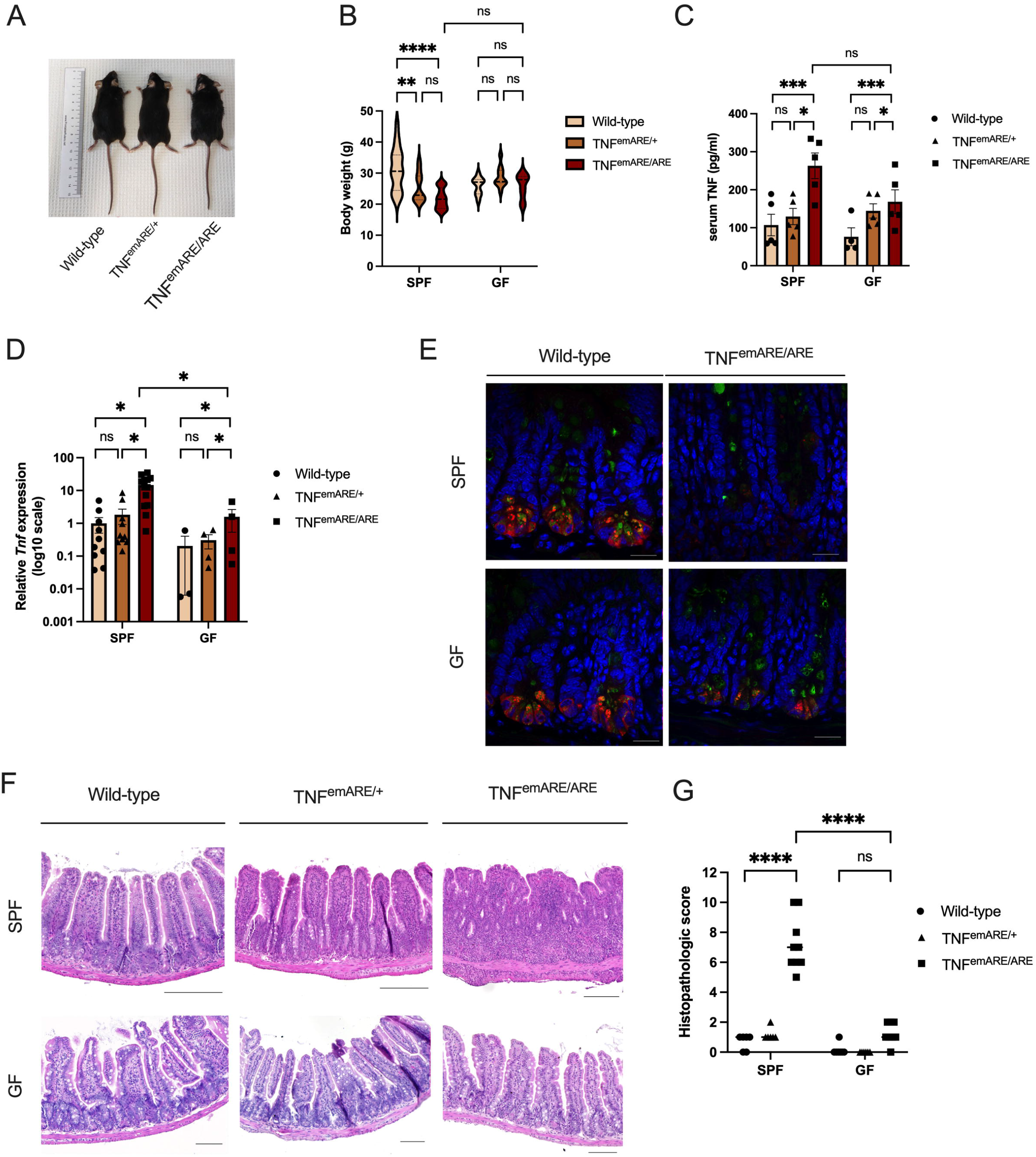
GF TNF^emARE^ mice are rescued from gut pathology. (A) Macroscopic picture of a GF male wild-type, male TNF^emARE/+^ and male TNF^emARE/ARE^ mouse shows no large differences in posture, except for a shorter tail of the homozygous mouse. Mice were between 20-24 w/o. (B) There is a trend in reduced body weight in 20-30 w/o SPF mice, which is less clear in the age-matched GF mice (SPF wild-type nF=6, nM=11; SPF TNF^emARE/+^ nF=12, nM=9; SPF TNF^emARE/ARE^ nF=16, nM=9; GF wild-type nF=5, nM=5; GF TNF^emARE/+^ nF=2, nM=8; GF TNF^emARE/ARE^ nF=3, nM=6). (C) Serum levels of TNF are elevated in hetero- and homozygous transgenic mice, in both SPF and GF conditions (n=5 mice/group). (D) Quantitative real-time PCR reveals increased expression of the *tnf* gene in the ileum in homozygous SPF and GF mice (SPF wild-type n=10, SPF TNF^emARE/+^ n=10, SPF TNF^emARE/ARE^ n=13; GF wild-type n=3, GF TNF^emARE/+^ n=4, GF TNF^emARE/ARE^ n=4). (E) Immunofluorescent ileal sections indicate Paneth cell depletion in SPF TNF^emARE/ARE^ mice, while these cells are still detected on ileal sections of GF TNF^emARE/ARE^ mice. Paneth cells = red (anti-lysozyme), mucins = green (WGA+UEA-1), nuclei = blue (Hoechst). (Scale bars: 100 μm). (F) Comparing SPF versus GF H&E sections of the ileum shows complete rescue of gut inflammation under axenic conditions in 20-30 w/o TNF^emARE/ARE^ mice. (G) Histopathological scoring quantitatively confirms rescue of ileal disease when mice are housed GF. Samples of 20-30 w/o mice were scored blindly. (SPF wild-type n=6, SPF TNF^emARE/+^ n=6, SPF TNF^emARE/ARE^ n=9; GF wild-type n=7, GF TNF^emARE/+^ n=6, GF TNF^emARE/ARE^ n=8) (Plots are represented as Mean +/- SEM)

Live *in-vivo* imaging using fluorodeoxyglucose (FDG) based PET-CT analysis was performed on SPF and GF raised wild-type and TNF^emARE/ARE^ mice, to visualize structural elements and sites of active inflammation in the whole body. PET-CT scans revealed multi-articular inflammation in front and hind paws, spinal column inflammation and non-congenital deformation (hyperkyphosis), in both SPF and GF raised TNF^emARE/ARE^ mice (Figure 3A). SPF TNF^emARE/ARE^ mice display elevated FDG signal in the abdominal area, indicating active intestinal inflammation. In contrast, no FDG signal is observed in the abdominal area of GF raised TNF^emARE/ARE^ mice (Figure 3A). We next performed H&E staining on histological sections of hind paws and found severe musculoskeletal pathology in both SPF and GF raised TNF^emARE/ARE^ mice, characterized by strong immune cell infiltration, bone erosion and bone marrow edema (Figure 3B). The histopathological scoring based on the calcaneocuboid joint, the calcaneus and the synovio-entheseal complex (Figure 3C) confirms similar musculoskeletal inflammation in colonized and axenic TNF^emARE/ARE^ mice. Also histological sections of the upper axial skeleton indicate spinal immune cell infiltration from the cervical to thoracic part, mainly along the spinal longitudinal ligaments and in the intravertebral discs of both colonized and axenic TNF^emARE/ARE^ mice (Figure 3D). In contrast to TNF^emARE/ARE^ mice, heterozygote TNF^emARE/+^ mice did not display clear inflammatory infiltrates in the peripheral joints or spinal deformations in either housing condition, in line with previous observations. To better characterize axial and peripheral pathology, μCT analysis was performed to visualize bone erosions in the calcaneus, and spondylosis and ankylosis of the vertebrae in detail. This high-resolution imaging technique is the preferred approach to study destructive effects of inflammation on the skeleton. Structural deformations (bone erosions) of the posterior calcaneus at the level of synovio-entheseal complex are clearly detectable, in both SPF and GF TNF^emARE/ARE^ mice (Figure 3E). Moreover, caudal vertebrae of the tail display vertebral bridging and fusion (Figure 3F).

**Figure 3:**
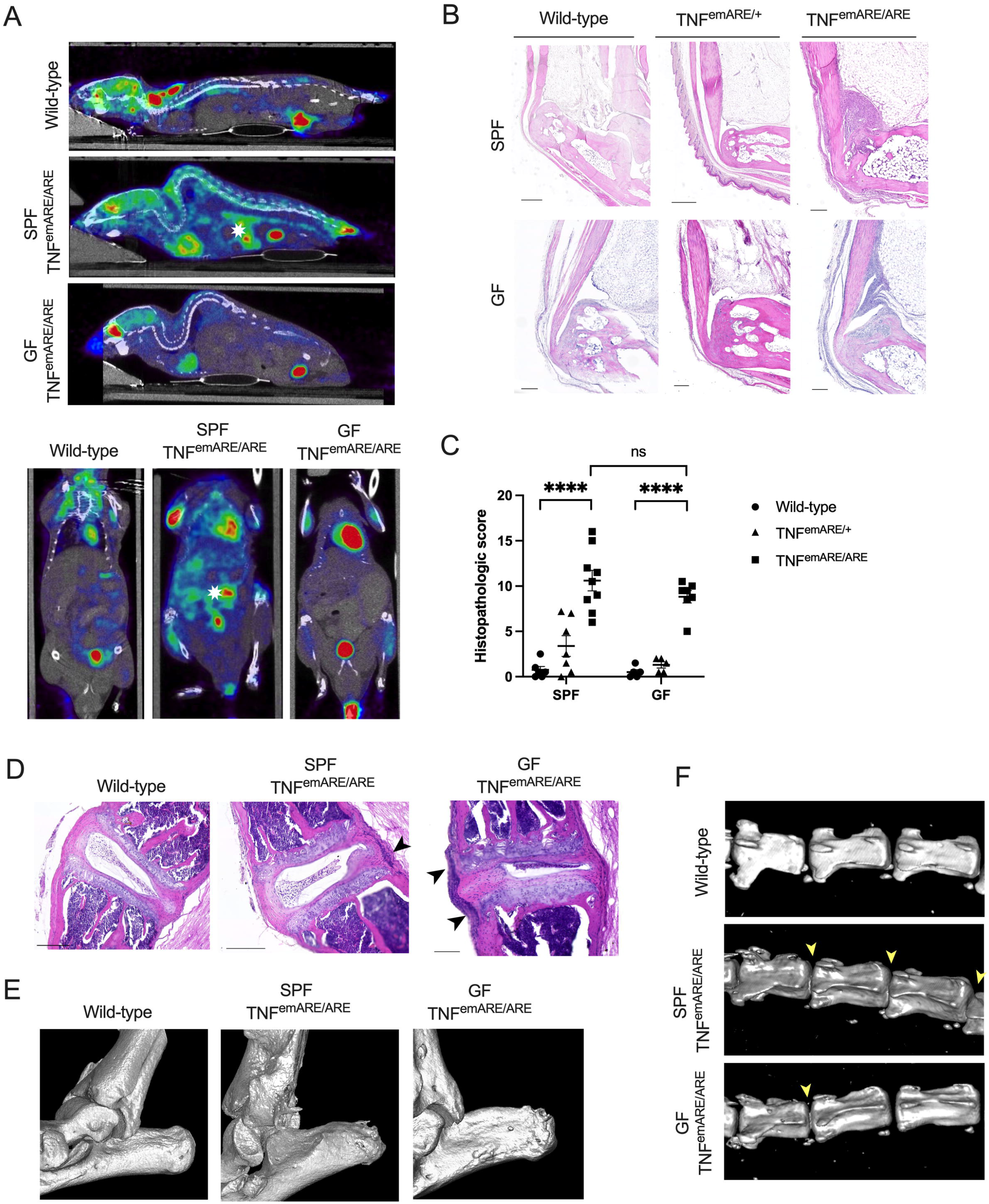
TNF^emARE/ARE^ mice display axial and peripheral arthritis in both SPF and GF conditions. (A) Wild-type and TNF^emARE/ARE^ mice housed in either SPF and/or GF conditions were subjected to PET-CT live *in vivo* imaging analysis. SPF TNF^emARE/ARE^ mice only show high uptake of FDG at the level of the intestines (indicated by a white asterisk), while SPF and GF TNF^emARE/ARE^ show inflammation at axial and peripheral joints and kyphosis of the spine. Regions in the mouse body showing high FDG uptake other than gut or joints are mainly artefacts because of tail vein injection, bladder content, physiologic myocardial uptake of FDG or active brown fat. (B) 20-30 w/o SPF TNF^emARE/ARE^ mice clearly suffer from inflammation in the synovio-entheseal complex, which is not rescued in age-matched GF mice. (Scale bars: 200 μm, except for SPF wild-type and SPF TNF^emARE/+^ 500 μm) (C) Histopathological scoring confirms severe arthritis development in both SPF and GF TNF^emARE/ARE^ mice. (SPF wild-type n=6, SPF TNF^emARE/+^ n=7, SPF TNF^emARE/ARE^ n=9; GF wild-type n=5, GF TNF^emARE/+^ n=5, GF TNF^emARE/ARE^ n=7) (D) Thoracic spinal histological H&E sections of an SPF wild-type versus SPF and GF TNF^emARE/ARE^ mouse reveal immune cells infiltrating along the spinal longitudinal ligaments (indicated by arrows). (Scale bars: 200 μm; except for GF TNF^emARE/ARE^ 100 μm) (E) μCT images of wild-type versus SPF and GF TNF^emARE/ARE^ mouse show structural deformations (bone erosions) of the posterior calcaneus in homozygous mice as a result of inflammation. (F) μCT images of tails of wild-type versus SPF and GF TNF^emARE/ARE^ mouse display vertebral bridging and fusion of sacral vertebrae in homozygous SPF and GF mice, as indicated by yellow arrows. All mice used for these experiments were between 20-30 weeks old. (Plots are represented as Mean +/- SEM)

To study the functional joint disability in TNF^emARE/ARE^ mice, we measured their grip strength compared to heterozygous TNF^emARE/+^ and wild-type littermates, both in mice raised in SPF and GF conditions. Both SPF and GF raised TNF^emARE/ARE^ mice have strongly reduced grip strength compared to wild-type controls (Figure 4A). Moreover, moving patterns of SPF and GF mice were studied using the CatWalk XT technology (Noldus). This technique allows to study mouse gait and locomotion in detail by capturing footprints as the mouse is traversing a glass walkway (representative videos in S4). Both SPF and GF TNF^emARE/ARE^ mice are characterized by aberrant gait, as they display smaller footprints, have lower foot surface pressure, and have reduced stride length compared to wild-type and TNF^emARE/+^ littermates (Figure 4B). Via illuminated footprint technology, each part of the paw that is in contact with the walkway can be detected and visualized. Footprints of both TNF^emARE/ARE^ SPF and GF mice show poorly defined foot contours (Figure 4C). These functional assays indicate reduced grip strength and abnormal gait as a result of severe arthritis in both SPF- and GF-raised TNF^emARE/ARE^ mice.

**Figure 4:**
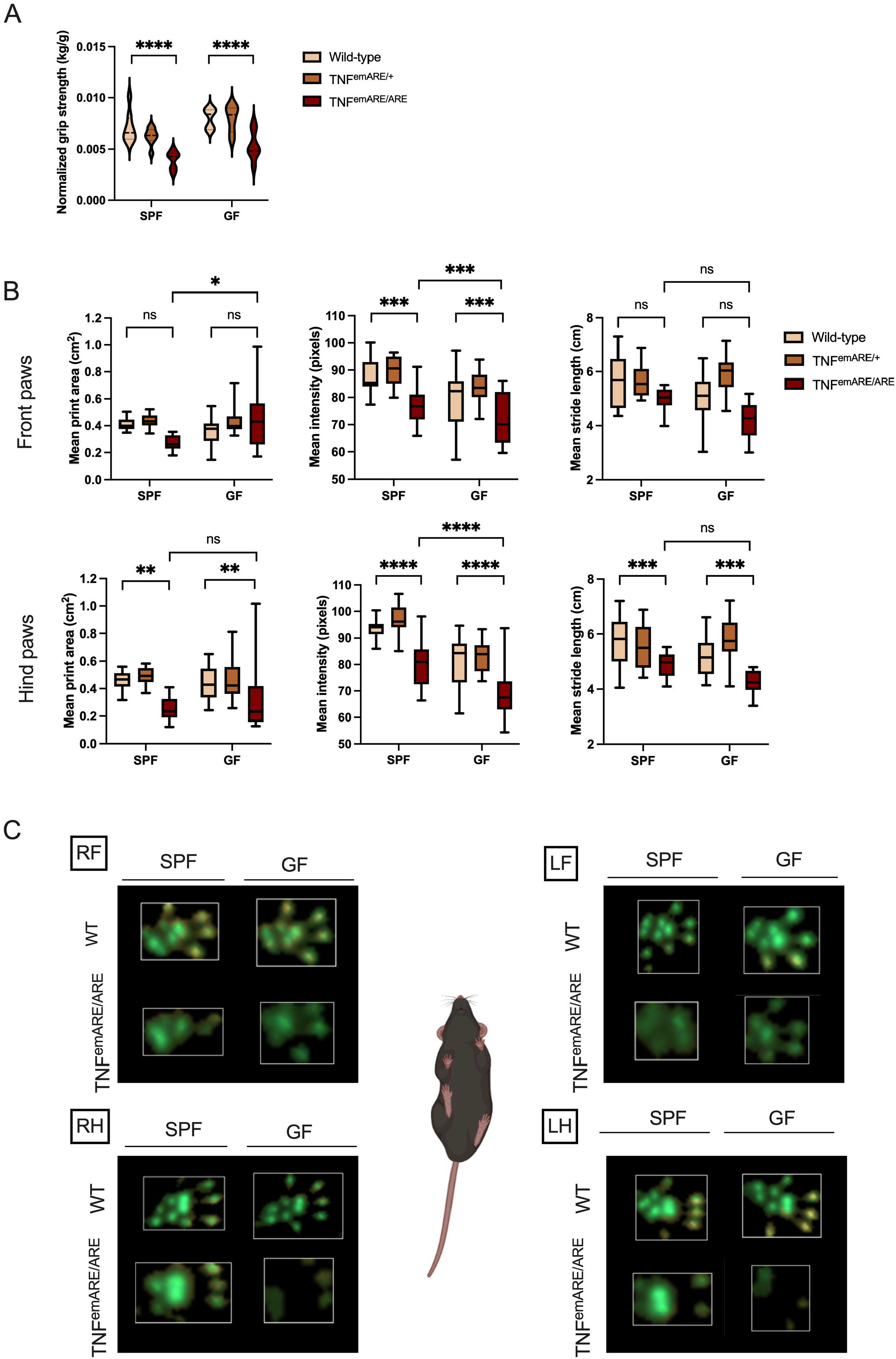
Gait analysis of TNF^emARE^ mice using the Noldus CatWalk gait analysis system. (A) Grip strength analysis of 4 paws of 15-30 w/o TNF^emARE^ mice shows a significant reduction in strength in TNF^emARE/ARE^ mice compared to their wild-type littermates, in both SPF and GF conditions. (SPF wild-type nF=4, nM=4; SPF TNF^emARE/+^ nF=4, nM=4; SPF TNF^emARE/ARE^ nF=4, nM=4; GF wild-type nF=4, nM=3; GF TNF^emARE/+^ nF=3, nM=4; SPF TNF^emARE/ARE^ nF=3, nM=4) (B) Gait analysis of TNF^emARE^ mice reveals distinct moving patterns of SPF and GF TNF^emARE/ARE^ mice. Homozygous mice tend to have smaller footprints, place their feet with lower mean intensity and generally have a smaller stride length. (SPF wild-type n=12, SPF TNF^emARE/+^ n=12, SPF TNF^emARE/ARE^ n=12; GF wild-type n=10, GF TNF^emARE/+^ n=20, GF TNF^emARE/ARE^ n=12) (C) Representative wild-type and TNF^emARE/ARE^ mouse footprints of mice raised under SPF versus GF conditions. Footprints of wild-type mice show precise prints visualizing the whole sole and toes, while in TNF^emARE/ARE^ mice the footprints are not clearly defined and prints of digits are missing. (RF = right front, LF = left front, RH = right hind, LH = left hind) Mice used for this analysis were between 20-40 weeks old. (Plots are represented as Mean +/- SEM)

Microbiota-derived succinate was previously shown to suppress ileal inflammation in TNF ^DARE^ mice, through tuft cell activation and expansion, while no data was shown on possible effects on joint inflammation (Banerjee *et al*., 2020). We investigated the gut-joint axis, and the assumption that joint inflammation is influenced by the intensity of intestinal pathology, by evaluating whether succinate supplementation via the drinking water (ad libitum) to SPF TNF^emARE/ARE^ mice improves not only gut but possibly also joint inflammation. We found succinate to promote tuft cell expansion and suppress ileal inflammation in TNF^emARE/ARE^ mice (Figure S5A, S5B). In contrast, succinate administration had no impact on musculoskeletal inflammation, as shown on histological sections of ankle joints (Figure S4C) and supportive quantitative scoring of disease severity (Figure S4D). These data show that improving gut inflammation does not necessarily improve joint inflammation, and rather support a functional disconnection of gut and joint pathology.

Together, these data clearly indicate that intestinal TNF-driven pathology in TNF^emARE/ARE^ mice is dependent on the intestinal microbiota, as GF TNF^emARE/ARE^ mice are fully protected from spontaneous ileitis development. In contrast, the microbiota is dispensable for TNF-driven musculoskeletal pathology in TNF^emARE/ARE^ mice, as GF TNF^emARE/ARE^ mice still develop severe arthritis. These data clearly indicate a functional disconnection of gut and musculoskeletal pathophysiology in this TNF-driven mouse model.

#### Joint inflammation in A20^myel-KO^ mice is driven by sterile triggers

In addition to a TNF-driven arthritis model, we also evaluated the importance of the microbiome for arthritis development in the IL-1β-dependent myeloid-specific A20 deficient (A20^myel-KO^) mouse model of arthritis (Matmati *et al*., 2011; Walle *et al*., 2014). A20^myel-KO^ mice were previously shown to develop spontaneous NLRP3 inflammasome- and IL-1β–dependent arthritis as a result of macrophage necroptosis (Matmati *et al*., 2011; Vereecke *et al*., 2014; Walle *et al*., 2014; Polykratis *et al*., 2019). However, A20^myel-KO^ mice do not develop intestinal pathology in small and large intestine, but are characterized by microbial dysbiosis (Matmati *et al*., 2011; Vereecke *et al*., 2014; Walle *et al*., 2014). IL-1*β* is known to play an important role in rheumatic diseases, but the upstream mechanisms leading to production of this interleukin are still incompletely understood (Lori Broderick, 2022). A20 negatively regulates inflammatory responses initiated by multiple pattern-recognition receptors (PRRs) and cytokine receptors. Moreover, polymorphisms in the *A20* locus are associated with many inflammatory and autoimmune diseases, including IBD, SLE and arthritis (Vereecke, Beyaert and van Loo, 2009; Ma and Malynn, 2012; Catrysse *et al*., 2014; Martens and van Loo, 2020). Transgenic A20^myel-KO^ mice were rederived in GF conditions by embryo transfer in the GF and gnotobiotic mouse facility at Ghent University. Since the phenotype of the A20^myel-KO^ model under SPF conditions has already been extensively studied and described (Matmati *et al*., 2011; Walle *et al*., 2014), we here merely focus on the comparison of arthritis features in A20^myel-KO^ mice housed under SPF versus axenic conditions.

Macroscopically, A20^myel-KO^ mice do not differ in body weight compared to their wild-type littermates, and this for both SPF and GF-raised mice (Figure 5A). However, macroscopic analysis reveals swollen ankle joints in SPF and GF A20^myel-KO^ mice (Figure 5B). To evaluate arthritis histologically, we performed H&E staining on foot sections and scored histopathological arthritis features by focusing on the calcaneocuboid joint, the calcaneus and the synovio-entheseal complex (Figure 5C, D). In 15 week old GF and SPF raised A20^myel-KO^ mice, immune cell infiltrates are clearly present in the fat pads and the synovium with occasionally affected cartilage and bone structures (Figure 5C). Remarkably, we observe massive bone marrow edema, which indicates severe systemic inflammation. In old GF A20^myel-KO^ mice (>30 weeks), the normal anatomical morphology of the hind paw is completely lost and we observe massive immune cell infiltration and loss of bone, cartilage and fat pads (Figure 5E). Similar to SPF raised A20^myel-KO^ mice (Matmati *et al*., 2011), GF A20^myel-KO^ mice show severe splenomegaly (Figure 5F), confirming a state of systemic inflammation. Together, these data indicate that also in this transgenic mouse model of innate (IL-1*β*)-driven arthritis, no causative role for the intestinal microbiota can be observed, as GF A20^myel-KO^ mice still develop severe arthritis.

**Figure 5:**
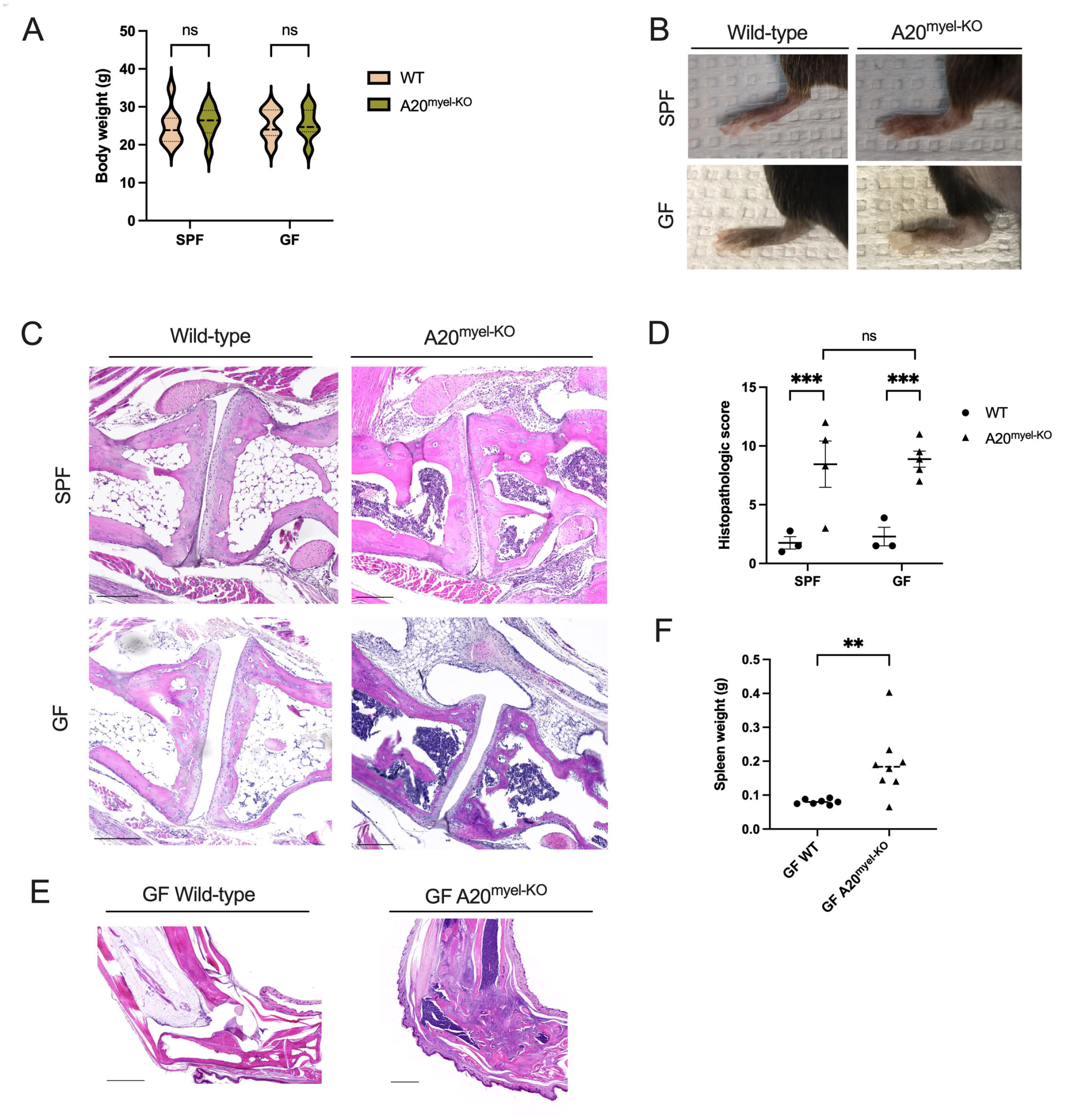
A20^myel-KO^ mice are not rescued from joint inflammation in axenic conditions. (A) Body weight between A20^myel-KO^ and wild-type mice does not differ significantly in both housing conditions. (SPF wild-type nF=5, nM=3; SPF A20^myel-KO^ nF=5, nM=3; GF wild-type nF=7, nM=7; SPF A20^myel-KO^ nF=4, nM=9) (B) Macroscopic images of hind paw of GF versus SPF A20^myel-KO^ mouse clearly show swelling of the ankle and toes. (Mice were 26-31 weeks old) (C) H&E sections of the calcaneo-cuboid joint of 5-15 w/o A20^myel-KO^ mice indicate immune cell infiltrates in the synovium and the fat pad and severe bone marrow edema. (Scale bars: 200 μm) (D) Histopathological scoring of joint inflammation of 5-15 w/o mice quantitatively confirms similar disease development in A20^myel-KO^ SPF and GF mice (SPF wild-type n=3, SPF A20^myel-KO^ n=4; GF wild-type n=3, GF A20^myel-KO^ n=5). (E) Histologic images of hind paws of old (44 w/o) GF wild-type (left) versus A20^myel-KO^ (right) mice. The morphology of the paw is completely lost in the transgenic mouse due to severe inflammation. (Scale bars: 1000 μm) (F) Splenomegaly in GF A20^myel-KO^ mice of 15 w/o indicates systemic inflammation in this mouse model (wild-type n=7, A20^myel-KO^ n=8). (Plots are represented as Mean +/- SEM)

In conclusion, our data demonstrate that in two genetic mouse models of arthritis, driven by either TNF or IL-1β, musculoskeletal disease develops independent of the microbiota, independent of gut inflammation, but instead is driven by joint-specific sterile factors.

## Discussion

Clinical observations in human IBD and arthritis suggest a common disease pathophysiology, commonly termed the ‘gut-joint’ axis. The fact that intestinal inflammation, though often subclinical, or an enteric infection, precedes joint inflammation in some patients, may support a causal relationship. The strongest genetic risk factor for SpA in humans is the human major histocompatibility complex (*MHC*) class I allele *HLA-B27*, which is believed to activate CD8+ cytotoxic T cells upon recognition of arthritogenic peptides. These arthritogenic peptides may be derived from the intestinal microbiota, and resemble self-peptides (molecular mimicry) which are subsequently targeted by effector T cells and cause joint inflammation (Pedersen and Maksymowych, 2019). Remarkably, HLA-B27 transgenic rats develop spontaneous gut and joint inflammation and are protected in GF conditions (Tautog *et al*., 1994). A recent study identified an arthritogenic strain of *Subdoligranum* that can drive systemic autoantibody production and joint inflammation in mono-associated mice (Meagan E. Chriswell, Adam R. Lefferts, Michael R. Clay, Alex Ren Hsu, Jennifer Seifert, Marie L. Feser, Cliff Rims, Michelle S. Bloom, Elizabeth A. Bemis, Sucai Liu, Megan D. Maerz,Daniel N. Frank, M. Kristen Demoruelle, Kevin D. Deane, Eddie A. James, Ja, 2022). Other experimental mouse arthritis models, including K/BxN, CIA and SKG require microbial triggers to drive type-3 immunity and arthritis development (Sakaguchi *et al*., 2003; Rehaume *et al*., 2014; Liu *et al*., 2016; Teng *et al*., 2017; Jubair *et al*., 2018). Furthermore, both IBD and SpA are characterized by changes in microbial community structure, composition and function, termed ‘dysbiosis’, which is suggested to prime mucosal immune cells which can traffic from the gut to the joints and cause inflammation (Gracey *et al*., 2020; Qaiyum, Lim and Inman, 2021). Photoactivatable transgenic Kaede and KikGR mice have provided evidence for trafficking of intestinal immune cells to extraintestinal sites including the joints, and gut-joint trafficking of colonic intraepithelial lymphocytes (IELs) was demonstrated in TNF^ΔARE^ mice (Morton *et al*., 2014; Adam R. Lefferts, Eric Norman, David J. Claypool, Uma Kantheti, 2022). Despite various experimental findings supporting a causal link between gut and joint inflammation, it remains unclear whether joint inflammation is critically dependent on microbial triggers or inflammatory cues from the gut. Its noteworthy that half of the SpA patients develop severe axial or peripheral joint inflammation in absence of subclinical gut inflammation. We therefore examined two independent transgenic mouse arthritis models, the TNF^emARE/ARE^ model which is driven by TNF, and the A20^myel-KO^ model which is driven by IL-1β. Our data demonstrate that severe arthritis can develop in both mouse models raised in GF conditions. In contrast, intestinal inflammation in TNF^emARE/ARE^ mice is fully microbiota-dependent. We hereby provide evidence that SpA-like arthritis in mice can be induced by sterile factors in absence of intestinal inflammation and the microbiota. Joint-associated sterile inflammatory triggers include various danger-associated molecular patters (DAMPs), of which the release is facilitated by mechanical loading, causing damage to the extracellular matrix or cell death in stromal and immune cells of the joint. These events promote recruitment of inflammatory immune cells and perpetuation of the inflammatory response through the production of inflammatory cytokines such as TNF, IL-6, IL1β, and IL-17. The role of mechanical loading in arthritis development was clearly shown in TNF^ΔARE^ mice, where hind limb unloading led to reduced entheseal inflammation and prevention of new bone formation (Jacques *et al*., 2014). IBD and SpA are complex multi-factorial diseases influenced by both environmental, genetic and immunological factors, with tightly intertwined inflammatory pathways underlying both diseases. We hypothesize that depending on the type and severity of the underlying genetic predisposition, depending on the microbiota composition, and depending on prior immune education, joint inflammation can be influenced to a varying degree by gut/microbiota derived mechanisms, but can also develop in response to sterile factors only.

## Materials and methods

### Generation of C57Bl6/J-Tnf^emARE1Irc^ mice

C57Bl6/J-Tnf^emARE1Irc^ (in this paper referred to as TNF^emARE^) mice were generated by the Transgenic Core Facility (TCF) of the VIB-Ugent Center for Inflammation Research (IRC). Guide sequences sequences 5’ GTGCAAATATAAATAGAGGG 3’ (sgRNA1) and 5’ GGAAGGCCGGGGTGTCCTGG 3’ (sgRNA2) were cloned in the BbsI site in the pX330 vector (addgene #42230). For sgRNA synthesis, the T7 promoter sequence was added to sgRNA forward primer and the IVT template generated by PCR amplification using forward primers 5’ TTAATACGACTCACTATAGGTGCAAATATAAATAGAGGG 3’ and 5’ TTAATACGACTCACTATAGGGAAGGCCGGGGTGTCCTGG 3’ for gRNA1 and gRNA2 respectively and reverse primer 5’ AAAAGCACCGACTCGGTGCC 3’. The T7-sgRNA PCR product was purified and used as the template for IVT using MEGAshortscript T7 kit (Thermofisher). Both sgRNAs were purified using the MEGAclear kit (Thermofisher). TNF^emARE^ mice were generated by injecting a mix of gRNA1 (10 ng/μl), gRNA2 (10 ng/μl), Cas9 protein (40 ng/μl; VIB Protein Service Facility) and Cas9 mRNA (20 ng/μl, Thermofisher) in C57BL/6J zygotes. Injected zygotes were incubated overnight in Embryomax KSOM medium (Merck, Millipore) in a CO2 incubator. The following day, 2-cell embryos were transferred to pseudopregnant B6CBAF1 foster mothers. The resulting pups were screened by PCR over the target region using primers 5’ TCTCATGCACCACCATCAA 3’ and 5’ GCAGAGGTTCAGTGATGTAG 3’. PCR bands were Sanger sequenced to identify the exact nature of the deletion. Mouse line TNF^emARE^ contains an allele with a deletion of 107 bp in the 3’ UTR of the Tnf gene at Chromosome 17:35418603-35418709 (GRCm39). This mutation is predicted to cause stabilization of the Tnf mRNA.

### Animal experiments

Tnf^emARE^ mice were generated as described above. Generation of A20^myel-KO^ mice was described previously (Vereecke *et al*., 2010; Matmati *et al*., 2011). Mice were housed in individually ventilated cages at the campus of UZ Ghent in a specific pathogen-free animal facility. Axenic mice were generated by embryo transfer in axenic recipients at the GF mouse facility of the University of Ghent. Axenic mice were housed under positive-pressure flexible film isolators (North Kent Plastics). All experiments were performed on mice of C57Bl/6 genetic background. All animal experiments were performed according to institutional (Ethical Committee for Animal Experimentation at Ghent University’s Faculty of Medicine and Health Sciences), national and European animal regulations.

#### Succinate experiment

Sodium succinate dibasic hexahydrate 99% (Sigma-Aldrich; S2378) was given to mice (n=8/genotype) of approximately 8 weeks old ad libitum via drinking water in a concentration of 120 mmol/L, for an average period of 15 weeks. Mice were followed-up by measuring body weight (every 2 weeks) and performing hemoccult fecal tests (week 10 and week 15). Endpoint of the experiment was after 15 weeks of treatment, mice were sacrificed and dissected.

### Grip strength test

The Bioseb Grip Strength Test was used to score functional disability in all 4 paws as well as the 2 front paws of mice. All scores were corrected for body weight.

### CatWalk gait analysis

Noldus CatWalk XT is a gait analysis system for rodents and was used to study differences in walking patterns between different mouse groups. Experiments were set-up with following parameters: minimum run duration of 0,5 seconds, maximum run duration of 5 seconds, maximum allowed speed variation at 60%.

Data were gathered and analyzed using the Noldus CatWalk XT software. Text on supplementary movies (genotype) was added using Descript 55.1.1 software.

### In-vivo imaging

#### PET-CT imaging was performed at the INFINITY lab of University Ghent

All animals were food deprived for at least 6 h prior to PET imaging. Mice were shortly anesthetized using a mixture of isoflurane and medical oxygen (5% induction, 1,5% maintenance, 0,3 l/min) to insert a catheter in one of the tail veins for tracer injection. Next, animals were intravenously injected with 10 MBq of FDG (Ghent University Hospital, Belgium) dissolved in 200 μl saline. Directly after tracer injection, the catheter was removed, mice were awakened and put into their cages. To reduce FDG uptake in brown fat, a heated blanket will be placed under the cage to keep the animals warm. In addition, the heated cage was placed in dark room to minimize tracer uptake into the Harderian glands. Forty minutes after tracer injection, the animals were placed under general anesthesia using an isoflurane mixture (5% induction, 1.5% maintenance, 0.3 l/min) and a 15-min total-body PET scan was acquired on a dedicated small animal PET scanner with sub-mm spatial resolution (B-Cube, Molecubes, Ghent, Belgium). Animals were placed in prone position, receiving further anesthesia through a nose cone. Body temperature was maintained at 37 °C by a heated bed. Each PET scan was followed by a total-body spiral high-resolution CT scan (X-Cube, Molecubes, Ghent, Belgium). The acquired PET data were iteratively reconstructed into a 192×192×384 matrix with 400μm isotropic voxel size. CT data were iteratively reconstructed into a 200×200×550 matrix with 200 μm isotropic voxel size. PET-CT images were processed and analyzed via Amide software(Loening and Gambhir, 2003).

### Ex-vivo imaging

Murine paws and spine were dissected, fixed in 4% formaldehyde for 48 hours and then kept in 70% EtOH. Ex-vivo high-resolution X-ray CT imaging of the murine hind paws and spines was performed at the Ghent University Centre for X-ray Tomography (UGCT). The samples were fixed in centrifuge tubes using wet cotton wool to avoid drying out of the samples during the μCT scan. The data was acquired using a commercial high-resolution CT scanner (CoreTom, TESCAN, Ghent, Belgium). For both paws and spines, 6 samples were imaged. The paws were imaged at a reconstructed voxel size of 7^3^ μm^3^, while for the spines a voxel size of 45^3^ μm^3^ was achieved. All samples are scanned using a tube voltage of 120kV and a hardware filter of 0.5 mm Al. Covering 360°, 2001 and 1501 projection images were acquired for the paws and spines, respectively, at an exposure time of 210 ms per projection image. After acquisition, tomographic reconstruction was performed using the proprietary software of the scanner system. The reconstructed volumes were exported as a stack of 16bit tiff slices, with the gray values representing the local reconstructed attenuation coefficients after rescaling which was fixed for each sample type (spine: −0,3 to 2 cm^−1^, hind paw: −1 to 2,2 cm^−1^).

μCT images have been processed using Fiji (Schindelin *et al*., 2012) and the MorpholibJ (Legland, Arganda-Carreras and Andrey, 2016) plugin before rendering them in 3D. A mask describing the region of interest (hind paw or axial skeleton) has been created by filtering the images using a gaussian blur with a sigma of 2 on the original image, setting a threshold on the image using the minimum method (Prewitt and Mendelsohn, 1966), filtering out the particles smaller than 50000 voxels and finally dilating the mask. The mask is then applied on the original image to remove the noise signal. The rendering in 3 dimensions has been done in Napari, a multi-dimensional image viewer for Python, using the iso-surface rendering.

### Histopathology

Murine gut sections were dissected and fixed in 4% formaldehyde for 24 hours. Paraffin sections were stained with Haematoxylin-Eosin (H&E) and ileal TNF^emARE^ sections were scored (blindly) by assessing villus architectural distortion (0-4), goblet cell depletion (0-4) and mononuclear cell infiltration (0-4) resulting in an overall score of 0-12. Images were acquired using Zeiss Axioscan and Zen Blue software.

Murine paws and spine were dissected, fixed in 4% formaldehyde for 48 hours and then decalcified using 5% formic acid for 8 consecutive days. Paraffin sections were stained with H&E for evaluation of inflammation and bone erosions. Disease development in hind paws was scored blindly by assessing the parameters in the table below, based on Yang-Hamilton (Yang and Hamilton, 2001) scoring and SKG scoring (Ruutu *et al*., 2012). Images were acquired using Zeiss Axioscan and Zen Blue software.

|  |  |
| --- | --- |
| Cuboidal joint: | - Immune infiltrates in synovium (0-2)<br>- Immune infiltrates in fat pad (0-2)<br>- Immune infiltrates in joint space (0-2)<br>- Immune infiltrates in cartilage (0-2) |
| Calcaneus: | - Bone erosion (0-2)<br>- Bone marrow edema (0-2) |
| Achilles tendon and Kargers' fat pad: | - Tendonitis (0-2)<br>- Immune infiltrates in synovium (0-2)<br>- Immune infiltrates in Kargers' fat pad (0-2)<br>- Immune infiltrates in joint space (0-2) |

### Immunofluorescent staining

Ileal sections were incubated in Dako antigen retrieval solution while being heated using a PickCell Electric Cooker. After cooling down, ileal sections were incubated with blocking buffer (goat serum 1/100) for 30 minutes at room temperature. Subsequently, sections were stained with primary antibody (rabbit anti-lysozyme (Dako; 1/500) or rabbit anti-DCAMKL1 (Abcam; 1/500)) overnight at 4C. After washing the slides with PBS, ileum sections were counterstained with DAPI (1/1000), UEA-1 Fluorescein (Vector laboratories; 1/1000) and WGA (ThermoFisher; 1/200) and incubated with secondary antibody goat anti-rabbit Alexa Fluor 568 (1/1000). After 1 hour, slides were mounted and later imaged with Zeiss AxioScan (10x) and processed with Zen Blue (anti-DCAMKL1 staining) or imaged with Zeiss LSM880 Airyscan (60x) and processed with Zen Black software (anti-lysozyme staining).

### Flow cytometry

Lamina propria isolation of small intestine samples of SPF TNF^emARE^ (wild-type n=5, TNF^emARE/+^ n=5, TNF^emARE/ARE^ n=5) mice was performed as described previously (Bain and Mowat, 2012). The isolated cells were used for extra- and intracellular staining for representative markers of the T cells and myeloid cells, in two separate panels (1×10^6^ cell/sample). Samples were analyzed using the five-laser BD LSRFortessa.

For the myeloid compartment, cells were stained with Fixable Viability Dye eFluor 506 (eBioscience; 1/300) for live/dead separation and only extracellular staining was done using following antibodies: CD19 (eBioscience; 1/400), CD3 (eBioscience; 1/200), NK1.1 (BioLegend; 1/200), anti-CD45 Alexa Fluor 700 (eBioscience 1/800), anti-Ly6G PercpCy5.5 (BD; 1/200), anti-Ly6C-APC (eBIoscience; 1/200), anti-Siglec F BUV395 (BD; 1/200), anti-CD11b BV506 (BD; 1/600), anti-CD64 BV711 (BioLegend; 1/100), anti-F4/80 Biotin (eBioscience 1/100), Streptavidin BV421 (BioLegend 1/1000), anti-CD11c PE-eFluor 610 (eBioscience; 1/300) anti MHC class II APC-eFLuor 780 (eBioscience; 1/800). For the T cell compartment, cells were stained with 7-AAD for live/dead separation, extracellular staining with anti-CD3 APC (eBioscience; 1/100), anti-CD4 APC Cy-7(BD; 1/200), and anti-CD8 V500 (BD; 1/100). Thereafter, cells were fixed and permeabilized using the Foxp3 Transcription Factor Staining Buffer Set (eBioscience; 00-5523-00). Finally, cells were stained intracellular with anti-Foxp3 Alexa Fluor 488 (eBioscience; 1/100) and anti-ROR t BV421 (BD; 1/100). Flow cytometry data were analyzed using FlowJo Software 10.8.1 and a sequential gating strategy. Regarding the tSNE plots, for the myeloid panel the analysis was performed on a total of 45000 CD45+ cells excluding the lineage (CD3, CD19, NK1.1) and for the T cell panel the analysis was performed on 173600 CD3+ cells. The cells were exported, concatinated and analyzed with FitSNE (Fast Fourrier Transform-accelarated Interpolation-based t-SNE) Flowjo plugin (version 0.5.1) (perplexity:20, Max iterations: 1000). Following dimensional reduction, coordinates for each t-SNE dimension (i.e., tSNE1 and tSNE2) in the two-dimensional plots were determined and integrated as novel parameters. For the myeloid panel, the bar plots showing percentages of immune cells represent proportions of every parent population. For the T cell panel, CD8+ and CD4+ cells are presented as percentages of the CD3+ parent population, CD4+RORgt+ and CD4+Foxp3+ cells are presented as percentages of the CD4+ population.

### ELISA

Blood was collected postmortem via cardiac puncture. Serum was isolated by 8 minutes centrifugation at 8000xg and stored at −20°C. Plates were coated with capture antibody anti-mouse/rat TNFa (eBioscience; 1/500) overnight at 4°C. Thereafter, plates were blocked with 0,1% casein blocking buffer for 2h by 27°C. Next, samples were added (undiluted) and incubation took 2h by 27°C. After washing, detection antibody anti-mouse/rat TNFa (eBioscience; 1/500) was added, followed by Avidin HRP enzyme (eBioscience; 1/100) and TMB substrate (BD Biosciences; 1/1). To end the reaction, stop solution (H2SO4, 1M) was added. Absorbance was read immediately at 450 nm. Concentrations were calculated in Graphpad Prism based on the standard curve.

### Quantitative real-time PCR (qPCR)

Tissue was lysed and homogenized using RLT and *β*—mercaptoethanol and the TissueLyser II (Qiagen). Total RNA was isolated using the RNeasy Mini Kit (Qiagen), according to the manufacturer’s instructions. The synthesis of cDNA was performed using QuantiTect® Reverse Transcription Kit (Qiagen), following the manufacturer’s protocol. For qPCR, SensiFAST SYBR NO-ROX (BioLine) and specific primers (*tnf* fwd TGTCTTTGAGATCCATGCCGT; *tnf* rev TCAAAATTCGAGTGACAAGCCTG) were used on LightCycler 480 (Roche). The reactions were performed in triplicates and the results were analyzed with qbase+ software. As housekeeping genes, GAPDH, Actb, Villin, Tbp, Ubc were used.

### Statistics

Statistical analysis was performed using analysis of variance (ANOVA) in Prism V9.2.0. Graphics were created using GraphPad Prism V9.2.0. Data are presented as mean ± SEM and p-values below 0,005 were considered statistically significant. Interpretation of asterisks on graphics:

|  |  |
| --- | --- |
| p-value > 0.05 | ns |
| p-value ≤ 0.05 | * |
| p-value ≤ 0.01 | ** |
| p-value ≤ 0.001 | *** |
| p-value ≤ 0.0001 | **** |

#### Body weight kinetic analysis

Body weights were analyzed as repeated measurements using the method of residual maximum likelihood (REML), as implemented in Genstat version 22. A linear mixed model (random terms underlined) of the form: body weight = constant + gender + genotype + time + gender×time + genotype×time + subject×time was fitted to the body weight data. The term subject×time represents the residual error term with dependent errors because the repeated measurements are taken in the same individual, causing correlations among observations. The uniform correlation structure was selected as best model fit based on the Akaike Information Coefficient. Times of measurement were set as equally spaced. The significance of the fixed main and interaction terms in the model, and of pairwise comparisons between genotypes across the time series, were assessed by an approximate F-test as implemented in Genstat version 22 (Gentstat.co.uk).

## Supporting information

supplemental movie GF TNFemARE

supplemental movie GFWT

supplemental movie SPF TNFemARE

supplemental movie SPF WT

## Supplementary material

Fig. S1 gives a schematic overview of the generation of a TNF-driven inflammation model (TNF^emARE^), Fig. S2 provides supportive data regarding the gut phenotype in TNF^emARE^ mice, Fig. S3 demonstrates TNF dependency of the inflammatory phenotype in TNF^emARE^ mice, Supplemental Movies S4 provide additional motion data on the CatWalk gait analysis experiments, Fig. S5 contains data on the succinate experiment using TNF^emARE^ mice.

## Acknowledgments

We would like to thank the HMI lab and the Dirk Elewaut lab for critical discussion and providing expert input, Kelly Lemeire from the IRC Immunohistochemistry core for assisting in staining protocols, the VIB Bioimaging Core Ghent for assisting in data acquisition and providing scripts to analyze microscopy data, the Animal House Facility staff in MRBII UZ Ghent for support with animal maintenance, the IRC Transgenic Core Facility for generating TNF^emARE^ mice and for rederiving TNF^emARE^ and A20^myel-KO^ mice germ-free, in collaboration with the Ghent germ-free and gnotobiotic mouse facility. We thank the VIB Flow Cytometry core facility for flow cytometry experiments and analyses. We thank Ghent University for funding this project (BOF-01N01019).

## Author contributions

L. Vereecke conceptualized and designed the experiments, wrote the manuscript, edited the final version and secured funding for the project; D. Elewaut co-funded the project, provided feedback on experimental results and edited the draft manuscript; A. Thiran designed and performed the experiments, analyzed data, designed figures, wrote the manuscript and edited the final version; I. Petta, designed and performed experiments and analyzed flow-cytometry data, G. Blancke, M. Thorp, M. Jans, T. Manuelo assisted in performing experiments; G. Planckaert assisted in performing experiments, analyzed data and provided expert input; V. Andries and K. Barbry were responsible for the GF facility and provided mice, T. Hochepied was responsible for the generation of transgenic TNF^emARE^ mice; C. Vanhove performed PET-CT imaging and associated data acquisition; I. Josipovic performed μCT imaging and associated data acquisition; E. Gracey provided expert input and assisted in analyzing data; E. Dumas conceptualized and designed the succinate experiments, G. van Loo generously provided A20^myel-KO^ mice for GF rederivation, and assisted in writing the final manuscript. All authors reviewed the manuscript.

## Abbreviations

CT: computed tomography
FDG: fluorodeoxyglucose
GF: germ-free
IBD: inflammatory bowel disease
μCT: micro-computed tomography
PET: positron emission tomography
RA: rheumatoid arthritis
SEC: synovial-entheseal complex
SpA: spondyloarthritis
SPF: specific-pathogen-free
w/o: weeks/old

## Legends of supplementary figures

**Figure S1. Generation of a TNF-driven inflammation model by targeting the AU-rich element of the *Tnf* gene.** (A) Schematic overview of a deletion of a 107bp fragment by CRISPR-Cas9 technology, followed by non-homologous end-joining (NHEJ). (B) A 107 bp fragment was deleted in the 3’UTR region (exon 4) of the *Tnf* gene on chromosome 17, exon 4. (C) Two guide RNA’s were used to specifically delete the targeted region.

*Figure created with Biorender.com*

**Figure S2: Gut phenotype of the TNF^emARE^ mouse model**. (A) Histological H&E sections of SPF TNF^emARE/ARE^ mice show an only minimally affected duodenum, while immune cell infiltrates are clearly detectable in jejunum. The ileum shows severe inflammation with loss of architecture. (Scale bars: 100 μm) (B) The colon of homozygous SPF mice remains healthy. (Scale bars: 100 μm)

**Figure S3: TNF^emARE/ARE^R1^-/-^ mice are rescued from gut and joint pathology**. TNF^emARE/ARE^R1^-/-^ mice are rescued and do not display ileal pathology nor arthritis. (Scale bars ileum: 200 μm, scale bars hind paws: 500 μm, scale bar focused images of SEC region: 200 μm)

**Supplemental Movies S4: SPF and GF TNF^emARE^ mice traverse the glass walkway of the CatWalk gait analysis system.** For every genotype, a representative video of a mouse traversing the glass walkway of the CatWalk gait analysis system (Noldus) is provided. Frame rate = 100 frames/second. Green intensity threshold = 0,10, amount of frames are displayed on every video. (RF = right front, LF = left front, RH = right hind, LH = left hind)

**Figure S5: Succinate supplementation induces tuft cell expansion and improvement of gut but not joint disease**. (A) Histological H&E sections of ileum indicate that succinate supplementation leads to improvement of ileal pathology. (Scale bars: 100 μm) (B) Immunofluorescent staining of DCLK-1 cells (red) shows expansion of tuft cells in succinate treated animals, in both wild-types and TNF^emARE/ARE^ mice. Mucins = green (WGA+UEA-1), nuclei = blue (Hoechst). (Scale bars: 100 μm) (C) Succinate treated mice are not rescued from arthritis development, disease severity is similar to control mice. (Scale bars: 500 μm) (D) Quantitative analysis of musculoskeletal pathology confirms no amelioration of joint disease in mice that received succinate treatment (Control groups n=4; succinate treated mice n=8).

