## Supplementary figures for "Sterile triggers drive joint inflammation in TNF and IL-1β dependent mouse arthritis models"

A

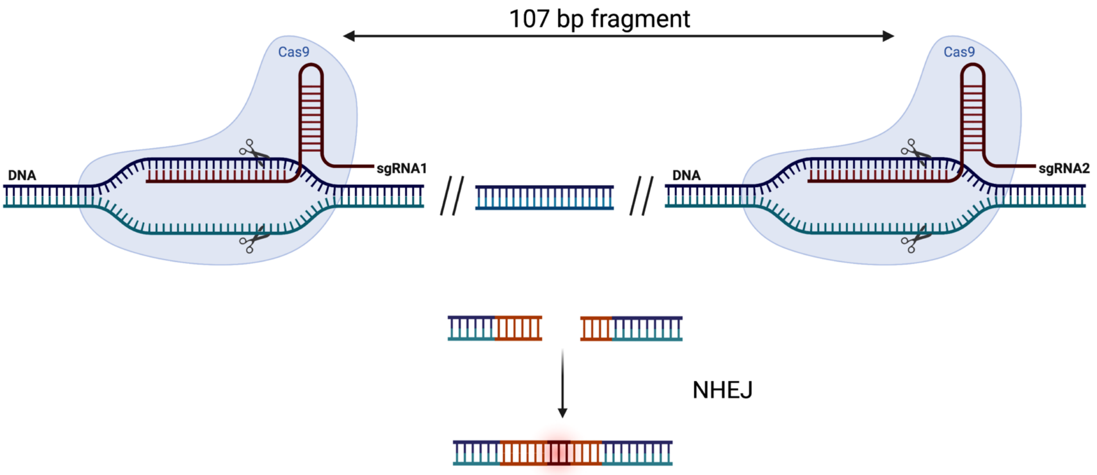


B

835

exon 4 CCTCACAGAG CCAG**CCCCCC TCTATTTATA TTTGCACTTA TTATTTATTA TTTATTTATT ATTTATTTAT**

941

exon 4 **TTGCTTATGA ATGTATTTAT TTGGAAGGCC GGGGTGTCCT GGAGGACCCA G**TGTGGGAAG CTGTCTTCAG

C

sgRNA1 5' GTGCAAATATAAATAGAGGG 3'
sgRNA2 5' GGAAGGCCGGGGTGTCCTGG 3'

Supplementary Figure 1

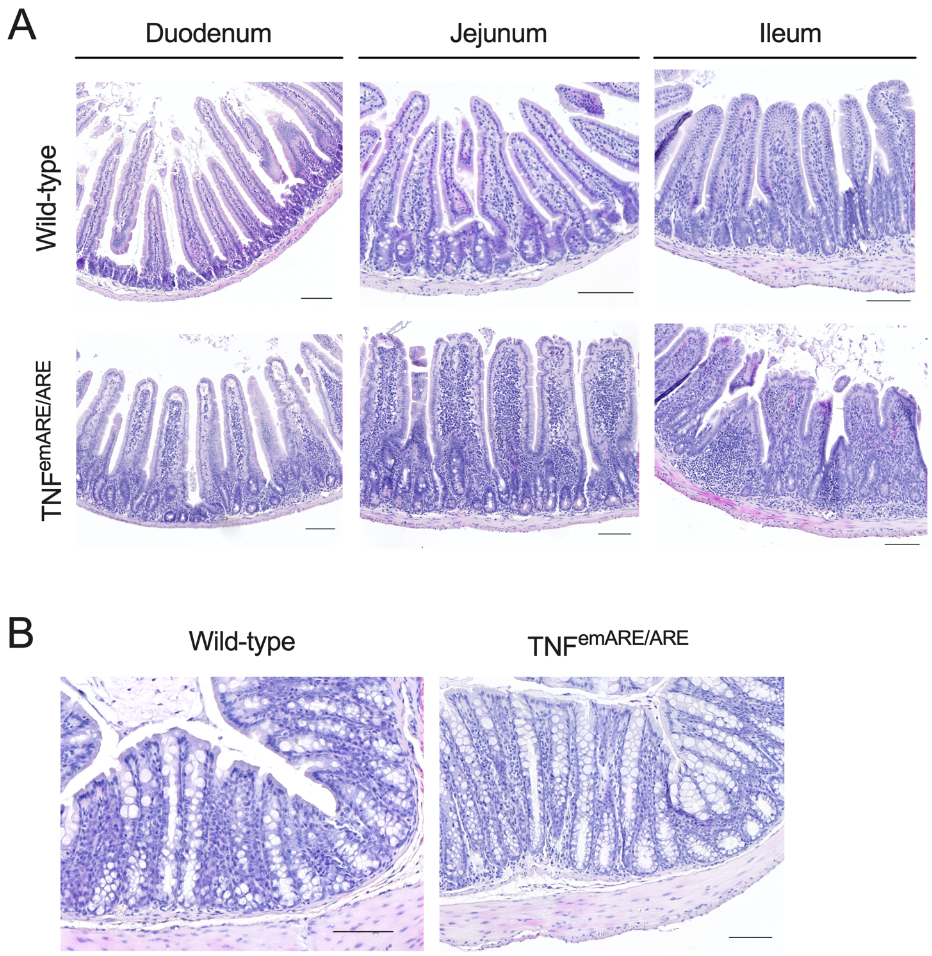
A
Duodenum
Jejunum
Ileum
Wild-type
TNF^emARE/ARE
B
Wild-type
TNF^emARE/ARE

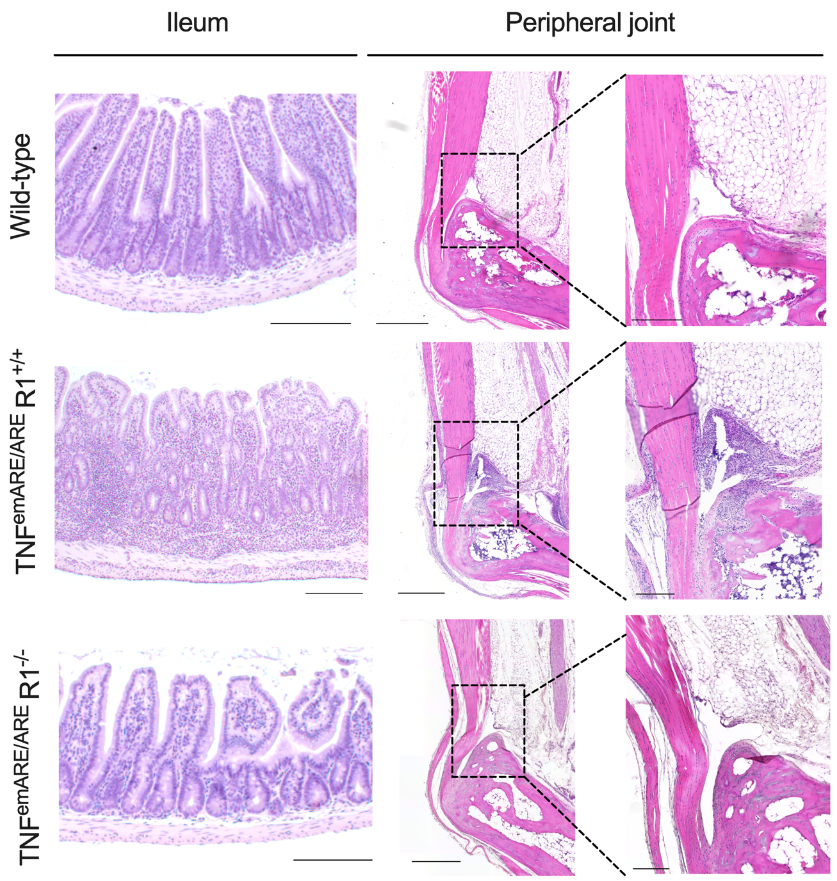
Ileum
Peripheral joint
Wild-type
TNF^emARE/ARE R1^+/+
TNF^emARE/ARE R1^-/-

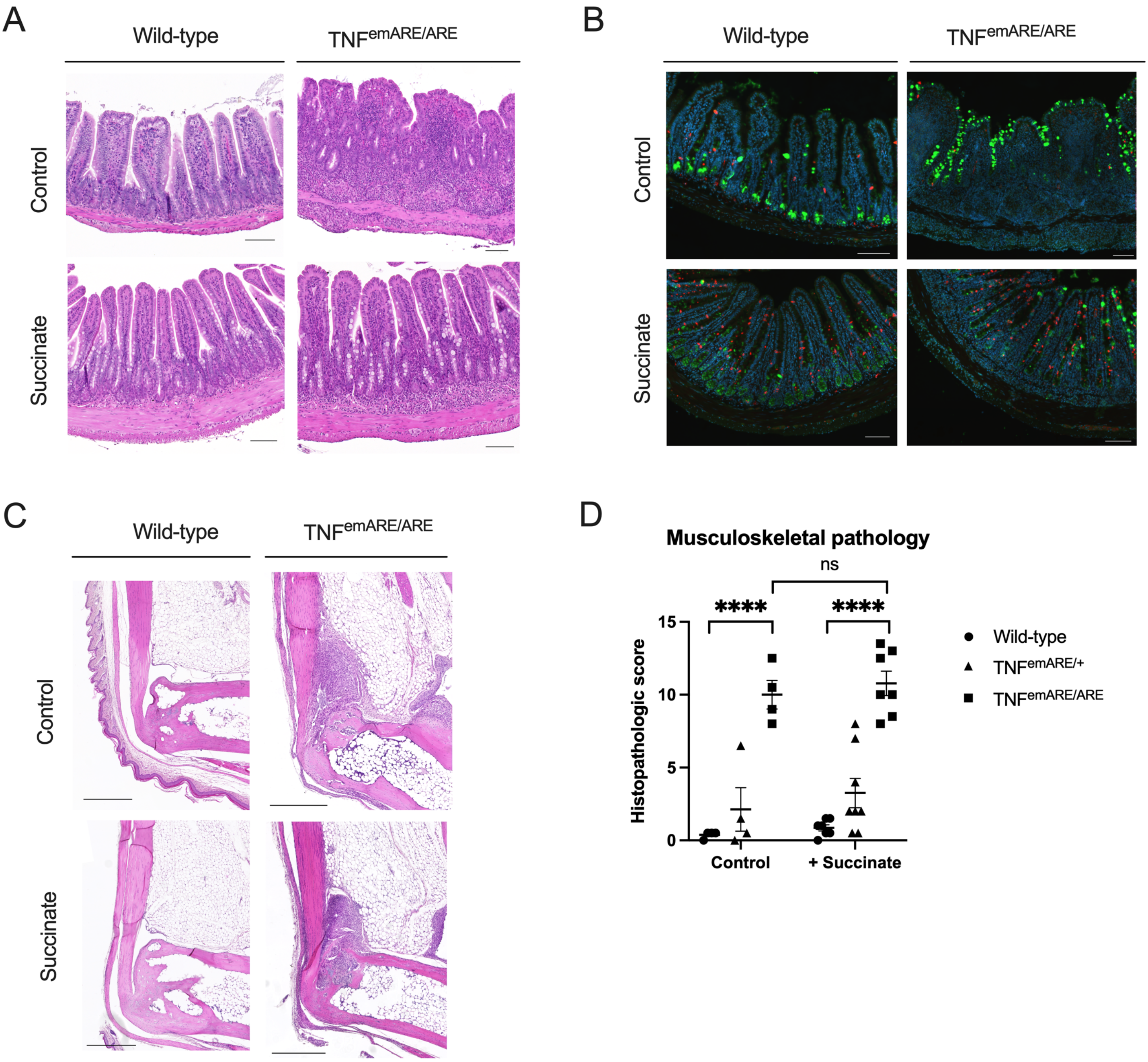
A
Wild-type
TNFemARE/ARE
Control
Succinate
B
Wild-type
TNFemARE/ARE
Control
Succinate
C
Wild-type
TNFemARE/ARE
Control
Succinate
D
Musculoskeletal pathology
ns
****
****
Histopathologic score
15
10
5
0
Control
+ Succinate
Wild-type
TNFemARE/+
TNFemARE/ARE
